# Interactions between non-structured domains of FG- and non-FG-nucleoporins coordinate the ordered assembly of the nuclear pore complex in mitosis

**DOI:** 10.1101/506469

**Authors:** Hide A. Konishi, Shige H. Yoshimura

**Affiliations:** Laboratory of Plasma Membrane and Nuclear Signaling, Graduate School of Biostudies, Kyoto University, Yoshida Konoe, Sakyo-ku, Kyoto 606-8501, Japan

**Author notes:** **Corresponding author:** Shige H. Yoshimura, Laboratory of Plasma Membrane and Nuclear Signaling, Graduate School of Biostudies, Kyoto University, Yoshida Konoe, Sakyo-ku, Kyoto 606-8501, Japan. Laboratory of Chromosome and Cell Biology, The Rockefeller University, New York, NY 10065, USA.

**Keywords:** Nuclear pore complex, Intrinsically disordered region, Molecular crowding, Self-assembly

## Abstract

In this study, we examined how channel-forming subunits of the nuclear pore complex (NPC) are assembled into a selective channel within a highly structured scaffold ring during post-mitotic assembly. We focused on non-structured domains of the scaffold Nups and performed *in vitro* self-assembled particle assays with those derived from channel-forming FG-Nups. We found that non-structured domains of ELYS and Nup35N interacted with channel-forming FG-Nups to form a self-assembled particle. Sequential addition of FG-Nups into the scaffold particle revealed that ELYS, which initiates post-mitotic NPC reassembly, interacts with early assembling FG-Nups (Nups98 and 153) but not middle stage-assembling FG-Nups (Nups58 and 62). Nup35, which assembles between the early and middle stages, facilitated the assembly of Nup62 into the early assembling Nups both *in vitro* and *in vivo.* These results demonstrate that ELYS and Nup35 have a role of facilitator in the ordered assembly of channel-forming FG-Nups during mitosis.

## Introduction

One of the largest protein complexes in eukaryotic cells is the nuclear pore complex (NPC), which spans the nuclear membrane and functions as the gate for macromolecular trafficking between the cytoplasm and nucleoplasm (1). The NPC is composed of several copies of more than 30 different nucleoporin (Nup) subunits (2). The NPC comprises several structurally distinct domains, such as the nuclear basket, cytoplasmic fibril, scaffold, and central channel, each of which is composed of defined sets of Nups. A number of studies using crystallography and electron tomography have revealed the molecular structures of the Nups and sub-complexes that comprise the scaffold (3). The entire architecture of the scaffold was then composited by integrating all available structural information for each of the subunits and sub-complexes with information regarding protein-protein interactions (4). Despite the low degree of amino acid conservation between Nups among different species, the entire architecture of the scaffold turned out to be well conserved. The outer ring containing the Nup107-160 sub-complex (Y-complex) is located at both the cytoplasmic and nucleoplasmic rims, and the inner ring, which contains the Nup93 sub-complex, is located deeper in the pore and connects the membrane and the central channel (3–5). Interactions between structured Nups and sub-complexes is thought to be stereospecific, resulting in a rigid architecture in the nuclear envelope.

In contrast to the scaffold, the central channel is formed primarily by a number of non-structured domains of Nups, which assemble via promiscuous interactions to form a highly crowded environment within the pore (6). Nups containing a large number of phenylalanine-glycine (FG) motifs (FG-Nups) in a long intrinsically disordered region (IDR-Nups) (7, 8) form a molecular sieve-like environment that functions as a selective barrier with respect to molecular transport. Small molecules (<40 kDa) diffuse passively, but large molecules (>40 kDa) are repelled unless they are carried by an appropriate transport receptor, such as a karyopherin (2, 9). Although several experimental and/or schematic models have been proposed to explain the structure and function of the barrier (10, 11), the molecular mechanism through which a large number of promiscuous interactions among IDR-Nups constructs a functional barrier within the rigid scaffold remains poorly understood.

Several lines of evidence have provided clues to enhance understanding of the mechanism of channel formation. Several FG-Nups self-assemble and form a hydrogel (10, 12) via hydrophobic interactions between Phe residues at a high protein concentration (≈10 mM) (13). A recent study reported that FG-domains of Nup98 spontaneously self-assemble into particles *in vitro* at physiologic concentrations (≈10 μM) (13). These findings suggest that the central channel of the NPC forms via self-assembly of non-structured FG-Nups. Another study using a crowding-sensitive fluorescent probe demonstrated that protein crowding is not homogeneous throughout the channel; “protein-rich” domains lie at both rims of the channel but not in the central cavity (6). This *in vivo* evidence suggests that even though FG-Nups self-assemble into protein-rich phases, several sets of Nups preferentially assemble into distinct protein-rich phases, possibly one at the cytoplasmic rim and another at the nucleoplasmic rim of the central channel. Furthermore, live-cell imaging of individual Nups during post-mitotic assembly of the NPC demonstrated that individual FG-Nups assemble orderly on the chromosome surface (6, 14, 15), suggesting that the assembly of FG-Nups is driven by a specific molecular mechanism and does not proceed solely via a random self-assembly–driven mechanism. Collectively, this evidence suggests that although FG-Nups self-assemble into protein-rich phases, a specific molecular mechanism regulates and/or coordinates the assembly of FG-Nups to construct distinct protein-rich phases within the central channel.

In this study, we focused on IDRs from FG-Nups in the central channel, as well as those from non-FG–Nups in the scaffold, and examined how they assemble and interact using an *in vitro* self-assembled particle assay. The results demonstrated that some non–FG-Nups, Nup35 and ELYS, interact with distinct sets of FG-Nups in a self-assembled particle and facilitate interactions with another set of FG-Nups. Our data suggest that IDRs in non–FG-Nups play an important role in the ordered assembly of FG-Nups to construct the functional selective barrier in the central channel of the NPC.

## Results

### Both FG-rich and non-FG IDRs form self-assembled particles in a crowded environment

Almost all FG-Nups contain a long stretch of IDRs (Figure S1A), each of which normally contains a number of FG motifs. In contrast, most non–FG-Nups do not carry IDRs, with the exception of Nup35, ELYS, and Tpr (Figure S1A). Nup35, which is found in the inner ring complex, contains a long IDR at its N-terminus (a.a. 1-160). ELYS, a component of the Y-complex in the outer ring, contains IDRs in the C-terminal half of the molecule (a.a. 1310-1559 and 1851-2275). Tpr is a component of the nuclear basket and has an extremely high number of IDRs throughout the molecule (especially in the C-terminal half). We focused on examining the interactions of these IDRs from non-FG-Nups, together with a number of FG-rich IDRs from FG-Nups (Nups214, 153, 98, 62, 58, and 50). IDR fragments examined in this study are summarized in Figure 1A and Table S1. Most IDRs from FG-Nups contain a large number of FG motifs, whereas Nup214-C and Nup50FG have only a few FG motifs and a relatively high content of charged residues (Figure 1A). The IDRs from non-FG-Nups have almost no FG motifs and are extremely rich in charged residues (Figure 1A). Purified IDR fragments were labeled with fluorescent dye, diluted into varying concentrations of polyethylene glycol (PEG)-containing buffer, and then observed under a fluorescence microscope. As summarized in Figures 1B-I, some of the IDR fragments self-assembled and form condensed particles in a PEG-dependent manner. Among FG-rich IDRs, Nups214FG, 153FG, 98FG, and 62FG self-assembled (Figures S1B-E), whereas Nups214-C, 58FG-N, 58FG-C, and 50FG did not (Figure S1F-I). The ability of forming particles appeared to correlate well with the FG content (Figure 1A). A FG-rich IDR from Nup214 (Nup214FG, containing 28 FG motifs) formed particles at a low PEG concentration, whereas Nup214-C (4 FG motifs) and Nup50FG (2 FG motifs) did not form particles even at a high PEG concentration (Figures 1D, S1H and S1I), suggesting that hydrophobic interactions are involved in particle formation by FG-rich IDRs (see later section).

**Figure 1.**
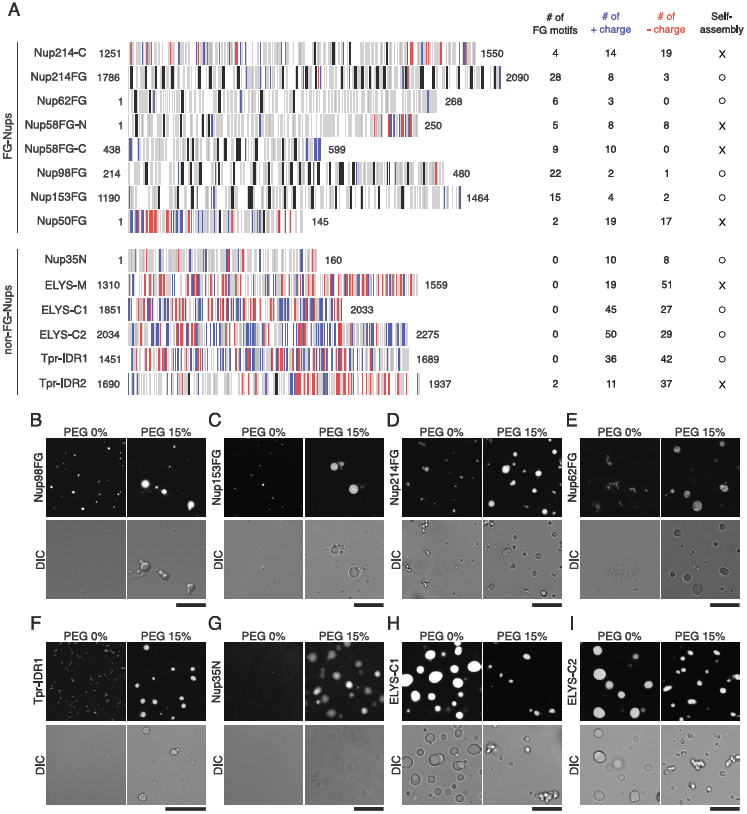
Self-assembly of IDR-Nups. (A) Amino acid compositions of IDR fragments used in this study. Vertical lines indicate the positions of positively charged (blue), negatively charged (red), and hydrophobic (gray) amino acids, and FG motifs (black). For visibility, other amino acid residues (polar and non-polar) are not shown here. Amino acid positions in the full-length protein are shown at both ends. The number of FG-motifs, positively and negatively charged residues, as well as the ability of self-assembly are shown on the right. (B-I) DIC and fluorescence images of the self-assembled particles of Nup98FG (B), Nup153FG (C), Nup214FG (D), Nup62FG (E), Tpr-IDR1 (F), Nup35N (G), ELYS-C1 (H), and ELYS-C2 (I) in the presence or absence of 15 % PEG3350. Scale bar: 10 μm.

Interestingly, some of the non-FG IDRs (Nup35N, ELYS-C1, -C2, and Tpr-IDR1) also self-assembled and form particles even though they do not contain FG motifs, but others did not (ELYS-M and Tpr-IDR2) (Figures 1F-I, S1J-M, -N, and -O). Amino acid sequence analyses revealed that all of these IDRs contained a number of charged residues, but only ELYS-M and Tpr-IDR2 had a large negative net charge (>25, Figure 1A and Table S1). This result suggests that particle formation by non-FG IDRs is driven predominantly by electrostatic interactions, but a large negative net charge may interfere with particle formation. Analysis of the secondary structure revealed that Tpr-IDR1, but not other IDRs, had a relatively high degree of coiled-coil structure (Figure S2B), although all of them had a high disorder probability. Circular dichroism (CD) spectra of Tpr-IDR1 indeed exhibited peaks at 222 nm and 207 nm, which indicates the presence of an α-helix (Figure S2C). These results suggest that coiled-coil interactions contribute to the self-assembly of Tpr-IDR1, as demonstrated in a previous study (16).

FG-rich IDRs in self-assembled particle exhibited a very slow exchange rate with the extra-particle environment and low mobility within the particle, as determined by fluorescence recovery after photobleaching (FRAP) analysis (Figures S2D-F) (13). In contrast, non-FG IDRs within particles exhibited faster recovery than FG-Nups (Figures S2G-I). This result is consistent with the previous report describing the difference between hydrophobic and electrostatic interaction in the self-assembly of proteins (17).

### Self-assembly of FG-rich IDRs and some non-FG IDRs is driven by hydrophobic interactions

We next characterized the protein-protein interactions that induce self-assembly of IDR-Nups by employing two reagents to quantify hydrophobic and electrostatic interactions: 1,6-hexanediol, an aliphatic alcohol that weakens hydrophobic interactions (11); and NaCl, which strengthens hydrophobic interactions but weakens electrostatic interactions. Self-assembly of FG-rich IDRs (Nup98FG and Nup214FG) was significantly inhibited with increasing concentrations of 1.6-hexanediol (Figures 2A, B and S3A), indicating the involvement of hydrophobic interactions in particle assembly. NaCl had less effect on FG-rich IDRs compared with 1,6-hexanediol; assembly of Nup214FG increased slightly (15-20%), whereas that of Nup98FG declined by 35-40% with increasing NaCl concentrations (Figures 2A, 2B, and S3B). These results suggest that the particle formation of FG-rich IDRs is facilitated by hydrophobic interaction.

**Figure 2.**
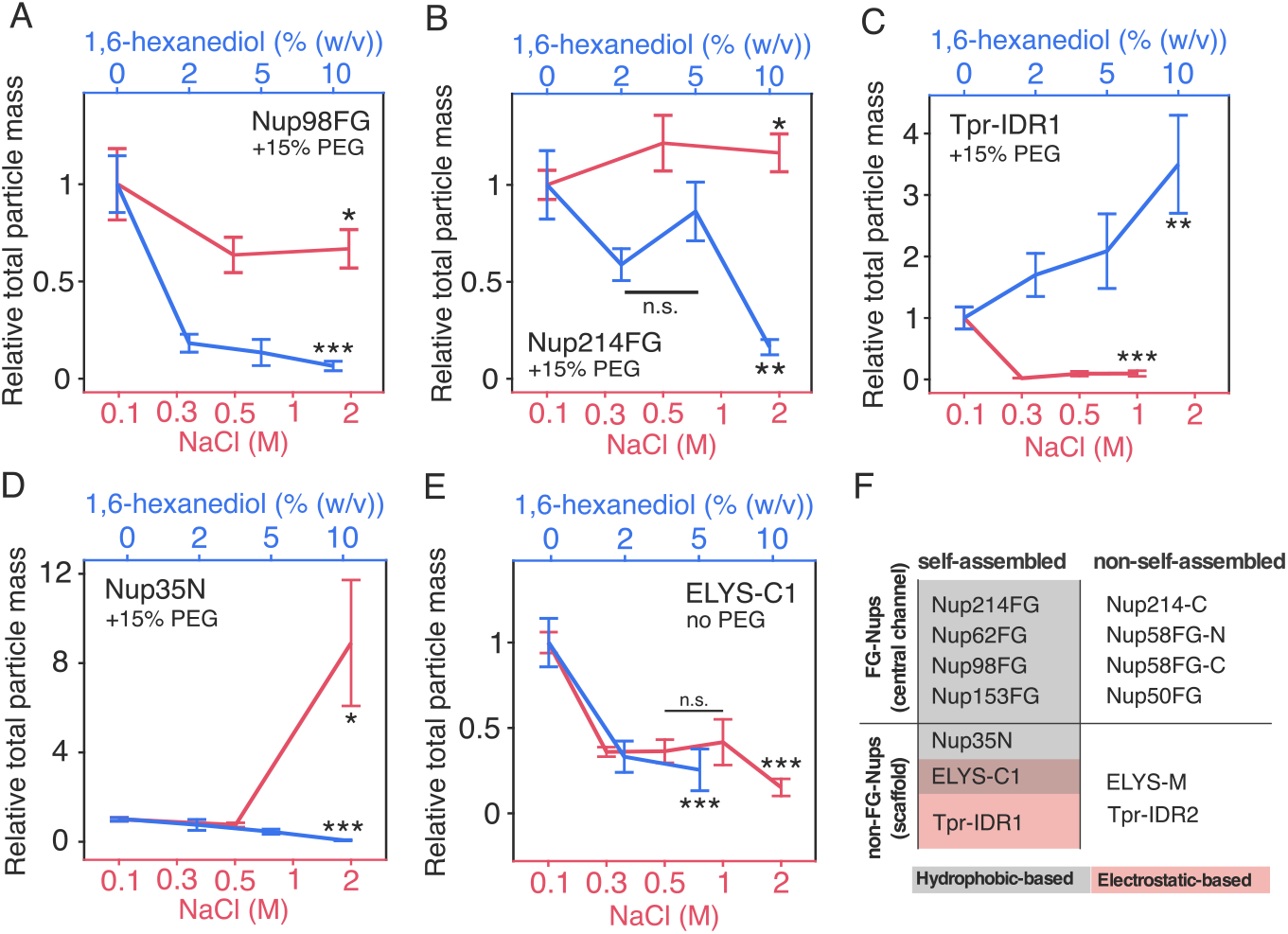
Self-assembly of IDR-Nups is driven by hydrophobic and/or electrostatic interactions (A-E) Co-line plot of NaCl- and 1,6-hexanediol-dependency in self-assembly of IDR-Nups (Nup98FG (A), Nup214FG (B), Tpr-IDR1 (C), Nup35N (D), ELYS-C1 (E)). Data are represented as mean ± SEM from three independent experiments. Each value is shown as a relative value to the standard condition (50 mM HEPES pH 7.4, 100 mM NaCl) in the presence or absence of 15 % PEG3350. (*: p<0.05, **: p<0.005, ***: p<0.0005). P-values were obtained by appropriate statistical tests (see Materials and Methods). (F) A summary of self-assembling properties of IDR-Nups. Hydrophobic-based (gray) and electrostatic-based (red) self-assembled IDR-Nups are highlighted.

In a clear contrast, the self-assembly of non-FG-Nup (Tpr-IDR1) was increased by 1.6-hexanediol but reduced with increasing NaCl concentrations (Figures 2C, S3A, and S3B), demonstrating that electrostatic interactions govern the self-assembly of Tpr-IDR1. Interestingly, Nup35N, which is also a non-FG IDR but with a small number of charged residues (Figure 1A) and high content of hydrophobic residues (Table S1), behaved as an FG-rich IDR; the particle mass increased as the NaCl concentration increased and declined as the 1,6-hexanediol concentration increased (Figures 2D, S3A, and S3B). Furthermore, ELYS-C1, a non-FG IDR containing a number of charged residues (Figure 1A), exhibited both electrostatic and hydrophobic characteristics (Figures 2E, S3A, and S3B). The assembly of ELYS-C1 was suppressed both by NaCl and 1.6-hexanediol, which could have been due to its unique amino acid sequence, which is characterized by short stretches (~20 amino acids) of charged and hydrophobic residues alternating with each other throughout the fragment (~200 amino acids) (Figure S3C). These results are summarized in Figure 2F. Most of the FG-rich IDRs self-assembled via hydrophobic interactions, but some non-FG IDRs (ELYS-C1 and Nup35N) also self-assembled via hydrophobic interactions.

### IDRs in ELYS and Nup35 interact with FG-IDRs

Two different IDRs, each labeled with a different fluorescent dye, were mixed in the same tube and observed under a fluorescence microscope to examine their interaction (Figure 3). One IDR was defined as the “host”, and the partition coefficient (PC) of the other IDR (client) in the host particle was then quantified (see Experimental Procedures). BSA was used as a standard of client. As summarized in Figure 3A and B, IDRs from FG-Nups generally co-localized well with FG-rich IDRs, although the individual PC value varied widely. FG-rich IDRs with a high FG content and a low content of charged residues (Nups214FG, 62FG, 98FG, and 153FG) exhibited high PC values, whereas IDRs with a low FG content and a high content of charged residues (Nups214-C, 58FG-C, and 50FG) exhibited low PC values, which are comparable to the standard BSA (Figures 3A and C).

**Figure 3.**
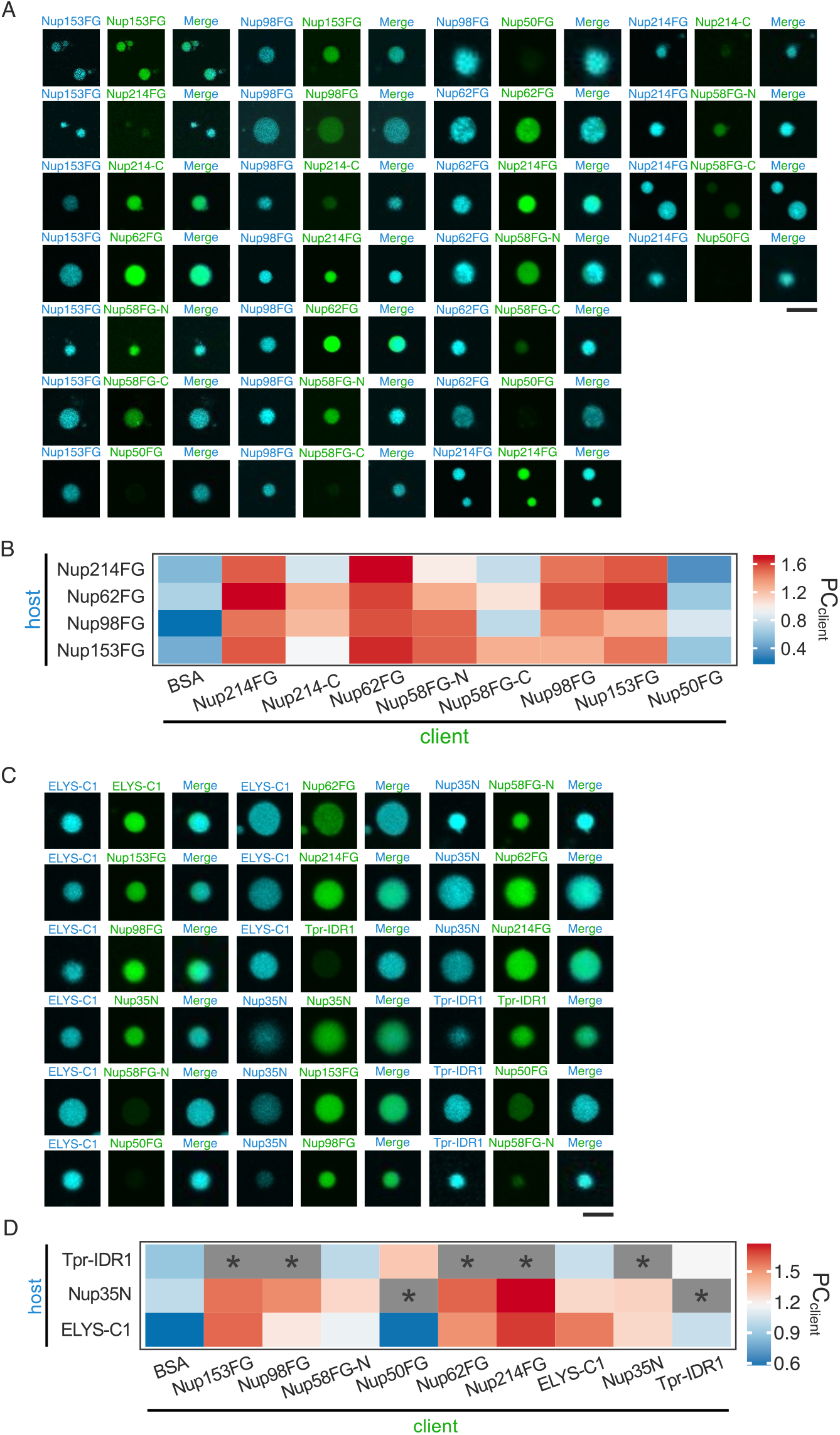
Interaction diagram of IDR-Nups in a self-assembled particle. Interaction between FG-Nups (A and B), and between FG-Nup and non-FG-Nup (C and D) were examined. (A and C) Fluorescence images of the self-assembled particle. Images were taken 5-10 min after mixing host and client proteins. Fluorescence signals of the host protein (labeled with ATTO390, left), and the client protein (labeled with ATTO488, middle), as well as the merged image, (right) are shown. Scale bar: 2 μm. (B and D) Tile plot of partition coefficient (PC) of client proteins against host proteins. Data are presented as an averaged value of the PC. An asterisk (*) indicates that client did not colocalize with host in the same particle as shown in Figure S3E. All of PC values are shown in Table S2.

The interactions between non-FG IDRs (ELYS-C1, Nup35N, and Tpr-IDR1) and between non-FG IDRs and FG-rich IDRs were also examined (Figures 3C and D). Tpr-IDR1, which self-assembled primarily via electrostatic interactions (Figure 2C), did not co-localize with most of the FG-rich IDRs except for Nup50FG (Figure 3C and D). In contrast, Nup35N, a non–FG-Nup that self-assembled via hydrophobic interactions (Figure 2D), co-localized with most of the FG-rich IDRs (Nups153FG, 98FG, 58FG-N, 62FG, and 214FG) (Figures 3C and D). Similarly, ELYS-C1 co-localized with FG-rich IDRs (Figures 3C and D). These results agree well with those shown in Figures 2D and E, showing that Nup35 and ELYS-C1, both of which are derived from non–FG-Nups, self-assembled via hydrophobic interactions, implying that Nup35 and ELYS-C1 play a role in the linkage between FG-Nups in the central channel and non–FG-Nups in the scaffold.

### ELYS-IDR interacts with early assembling but not middle stage–assembling FG-Nups

In mitosis, the NPC disassembles during prophase and reassembles during anaphase via the ordered assembly of individual Nups. ELYS has been shown to play a critical role in the initial step of reconstruction by binding to the chromosome surface (18). Nups located on the nucleoplasmic side of the central channel (e.g., Nups153 and 98) and some of the scaffold Nups (e.g., Nups133 and 160) (early assembling Nups) follow ELYS. Nups in the inner ring complex (Nups35 and 155) and membrane-spanning Nups (Pom121 and Ndc1) then assemble, which is followed by Nups in the central cavity (Nups54, 58, and 62) (14, 15) (Figure 4A). To investigate how the ordered assembly of channel-forming FG-Nups is regulated, we sequentially added FG-rich IDRs as clients and examined how they are incorporated into the ELYS-C1 particle as a host (Figure 4B, upper panel).

**Figure 4.**
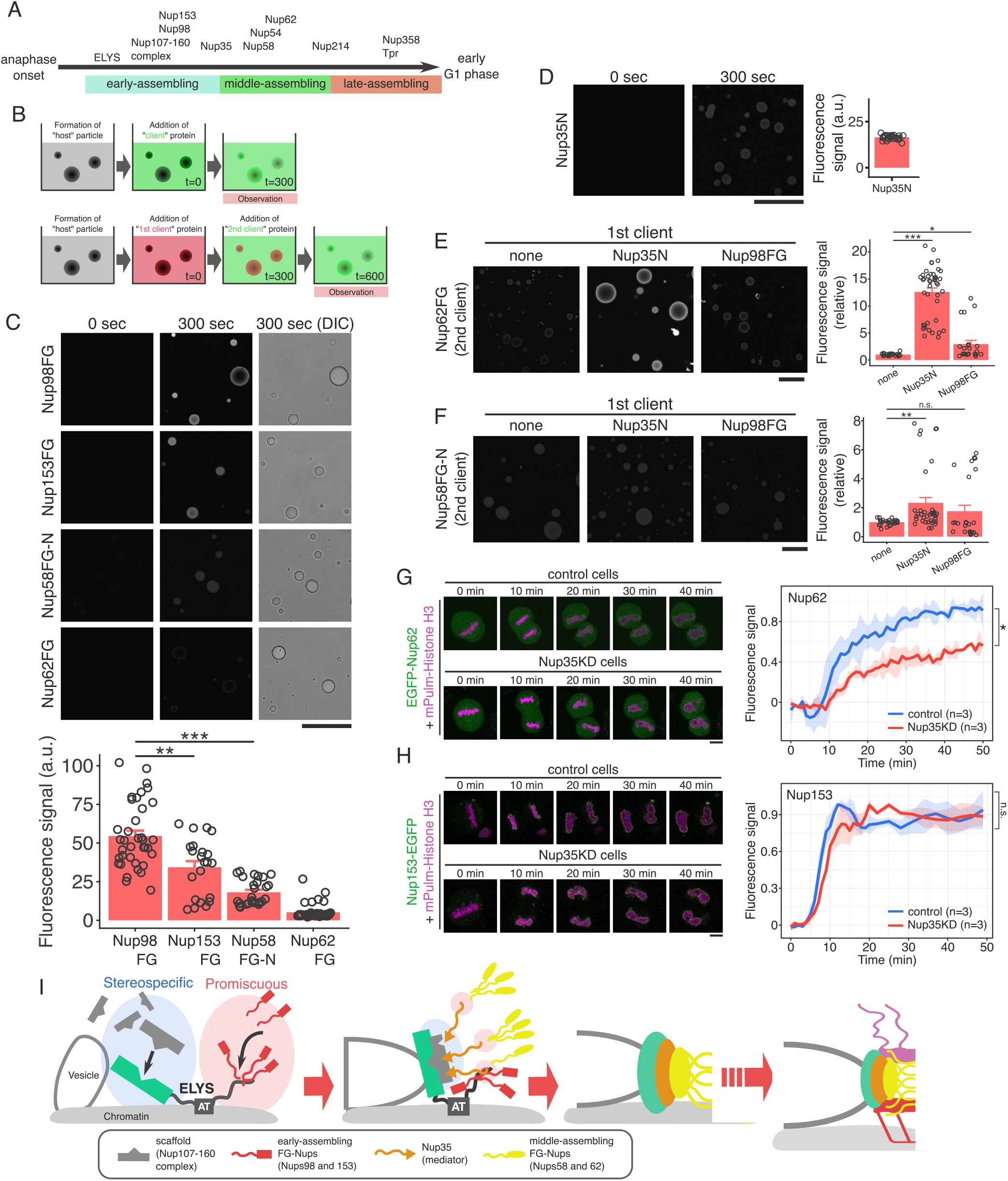
Nup35 facilitates the assembly of middle-assembling FG-Nups during the post-mitotic NPC reassembly. (A) The order of Nup reassembly in the post-mitotic NPC reassembly. Early-, middle-, and late-stage assembling Nups are plotted along the time after anaphase onset. (B) Experimental schemes of the *in vitro* sequential addition assay. A fluorescently-labeled client protein was added to the host particle (ELYS-C1) solution and the fluorescence signal was observed under the microscope 300 sec after the addition (upper panel). Alternatively, a fluorescently-labeled 1st client was first added to the host particle solution (t = 0), and then the 2nd client, which was labeled with different fluorescent dye, was added 300 sec after the addition of the 1st client (t = 300). The fluorescence signal of the 2nd client was observed under the microscope 300 sec after the addition of the 2nd client (t = 600) (lower panel). (C) A summary of sequential addition assay as depicted in the upper panel of (B). DIC image of the particle (300 sec after the addition) and the fluorescence images of the client (Nups98FG, 153FG, 58FG-N, and 62FG) (0 and 300 sec) are shown (upper panels). The fluorescence intensity of the client in the host particle was quantified and plotted in the bottom panel. Data are represented as mean ± SEM from three independent experiments. Multi-color images of both the host and the clients are shown in Figure S4A. Scale bar: 10 μm. (D) Fluorescence images of Nup35N, which had been incorporated into the host particle (ELYS-C1). The fluorescence intensity was quantified as in (C), and plotted in the right panel. (E and F) Fluorescence images of the 2nd client proteins in the sequential addition assay. As described in B (bottom panel), the 1st client (Nup35N or Nup98FG) was first added to the host particle (ELYS-C1) (t =0) (images are shown in Figure S4B), and then the 2nd client (Nup62FG (E), Nup58FG-N (F)) was added (t = 300). Images of the 2nd client were taken 300 sec after the addition (t =600), and are shown in (B, lower panel). The fluorescence intensity of the 2nd client protein (Nup62FG or Nup58FG-N) was quantified and plotted as a relative value to the value without the 1st client (none). Data are represented as mean ± SEM from three independent experiments. Scale bar: 10 μm. *: p<0.05, **: p<0.005, ***: p<0.0005, n.s.: no significance. P-values were obtained by appropriate statistical tests (see Materials and Methods). (G and H) The effect of Nup35-knockdown on the post-mitotic assembly of Nup153 and Nup62. EGFP-fused Nup62 (G) or Nup153 (H) was expressed in Nup35-knocked-down HeLa cells, together with mPlum-fused histone H3, and was subjected to the time-lapse imaging during mitosis. Fluorescence signal at the chromosome rim was quantified as described in Materials and Methods, and presented as relative values to the maximum value. Data are shown as mean ± SEM. The number of the analyzed cells is also indicated. *: p<0.05 and n.s.: no significance. P-values were obtained by appropriate statistical tests (see Materials and Methods). Scale bar: 10 μm. (I) Schematic model of the post-mitotic NPC reassembly. See text for detail.

IDRs from early-assembling FG-Nups (Nups98FG and 153FG) were rapidly incorporated into the ELYS-C1 particle after addition (Figures 4C and S4A), consistent with the results shown in Figure 3D and F. Interestingly, lower amounts of 58FG-N (which assembles during the middle stage) were incorporated into the ELYS particle, compared with the early assembling Nups (Figure 4C). Furthermore, Nup62FG (which assembles during the middle to late stages) was not incorporated into the ELYS particle (Figure 4C), despite generally co-localizing in the same particle when mixed (Figures 3D and F). These results demonstrate a clear correlation between the interaction between ELYS particle and the order of the FG-Nups during post-mitotic reassembly; ELYS, which binds to the chromosome surface at the beginning of NPC reassembly, readily recruits Nups98 and 153 (early assembling Nups) but not Nups58 and 62 (middle stage–assembling Nups).

### Nup35 facilitates the assembly of middle stage–assembling Nups

In addition to ELYS, Nup35N also interacted with FG-rich IDRs in our particle assay (Figure 3). As Nup35 appears on the chromosome surface during the early to middle stages of post-mitotic reassembly (19) (Figure 4A) and interacts with the structured domain of Nup62 (4), it can be hypothesized that it facilitates the assembly of middle stage–assembling FG-Nups. We therefore tested the possibility of Nup35N to facilitate the incorporation of middle stage–assembling Nups into ELYS particles. Nup35N, as 1st client, was pre-incorporated into the ELYS-C1 particle in part to mimic the early to middle stages of post-mitotic reassembly (Figure 4B lower panel). As shown in Figure 4D, Nup35N was incorporated into the ELYS-C1 particle at a rate comparable to that of other early assembling Nups (Nups153FG and 98FG) (Figure 4C), consistent with results shown in Figures 3D and F. The middle stage–assembling Nup58 or 62 was then added, and its incorporated fluorescence signal in the particle was quantified. Interestingly, incorporation of Nup62FG into the ELYS-C1 particle was enhanced when the particle contained Nup35N (Figure 4E). A similar effect of Nup35N was observed with the incorporation of Nup58FG, although the effect was smaller (Figure 4F). It should be noted that no such stimulatory effect was observed with Nup98FG, an early assembling FG-Nup; pre-incorporation of Nup98FG into ELYS-C1 particle did not facilitate the incorporation of Nups62 and 58 (Figures 4E and F). These results demonstrate that Nup35N facilitates the interaction of middle stage–assembling Nups with early assembling Nups.

Finally, we examined the role of Nup35 in the post-mitotic NPC reassembly process *in vivo.* An EGFP-fused FG-Nup (Nup62 or 153) was expressed in Nup35 (or control) knock-down (KD) cells, and the appearance of fluorescence signals on the chromosome rim during telophase and anaphase was observed by time-lapse imaging. Signal associated with Nup62 began to appear on the chromosome surface ~10 min after anaphase onset and gradually increased for 30 min until it reached a plateau (Figure 4G). In contrast, Nup35-KD cells exhibited a significant delay in Nup62 assembly; it began to appear 10 min after the anaphase onset, which is similar to control knock down, increased slowly, and did not reach the plateau even after 50 min. It should be noted that the effect of Nup35 KD was not observed in the assembly of Nup153, an early assembling Nup; it began to appear 3-5 min after the anaphase onset, and reached the plateau at 12-15 min in both control and Nup35-KD cells (Figure 4H). These results provide *in vivo* evidence that Nup35 facilitates the assembly of middle stage–assembling Nups during post-mitotic NPC reassembly.

## Discussion

In this study, we investigated the self-assembling properties of IDRs derived from FG-Nups and non-FG-Nups as well as their interaction patterns using an *in vitro* self-assembled particle assay. The results demonstrated that i) many FG-rich IDRs self-assemble via hydrophobic interactions and co-assemble (co-localize) well in the same particle; ii) some IDRs from non–FG-Nups self-assemble via electrostatic interactions, whereas others (Nup35N and ELYS-C1) assemble via hydrophobic interactions and interact with FG-rich IDRs; iii) The IDR of ELYS (ELYS-C1) interacts with early assembling FG-rich IDRs (Nups98FG and 153FG) but not with middle stage–assembling Nups (Nups58 and 62); and iv) Nup35N facilitates the assembly of middle stage–assembling FG-rich IDRs (Nups58FG and 62FG) into the ELYS particle. These results demonstrate the important roles of IDRs from scaffold Nups in coordinating the assembly of FG-Nups. It can be hypothesized that although self-assembly of Nups is generally driven via promiscuous interactions among IDRs, “ordered” assembly of Nups and the construction of a functional selective channel during the post-mitotic reassembly process could be mediated by a few “facilitator subunits,” such as ELYS and Nup35. Functional analyses of these mediator subunits could elucidate the mechanism underlying the construction of the central channel, and together with structural information regarding the scaffold, such analyses could elucidate the relationship between the structure and function of the mitotically formed NPC.

The molecular architecture of the scaffold has been studied extensively using crystallography, electron tomography, and mass spectroscopy (3, 20). Nup35 is a component of the Nup93 complex, which is the major structural unit of the inner ring, and interacts with Nups in the same complex (Nups93, 155, and 205) as well as other Nups (Nup98 and Ndc1) and the membrane (4). A recent study reported the crystal structure of the Nup93 complex with structured domains of the central channel components Nups62, 58, and 54 (21), demonstrating that the Nup93 complex links the scaffold and central channel. Although the entire structure of Nup35 was not elucidated in the crystallographic study, it is possible that the IDR of Nup35 associates with the IDRs of Nups62, 54, and 58. Our *in vitro* self-assemble particle assay indeed demonstrated that Nup35N, which has no FG motif, interacted with most of the FG-rich IDRs examined (Figure 3D and F).

From a structural perspective, the interaction between Nup35N and FG-rich IDRs could be attributed to its unique amino acid composition, which is rich in proline (≈20%) and glycine (≈11%) (Figure S3D). A large number of proline residues could aid the formation of self-assembling particles, as well as hydrophobic interactions with FG-rich IDRs. It was previously demonstrated that interactions between proline-rich motifs in N-WASP and the SH3 domain of a partner molecule are important for multivalent interactions between them (22). Proline is also known to break secondary structures (a-helixes and ß-sheets) (23–25) and confer a high degree of flexibility on polypeptides. Such flexibility could increase the mobility of molecules within the particle, thus facilitating the incorporation of other molecules into the host particle.

In addition to Nup35, a fragment of the ELYS IDR (ELYS-C1) also interacted with FG-rich IDRs, indicating that it can also function as a facilitator between the scaffold and central channel. ELYS contains several distinct functional domains in addition to its IDR and was shown to play an important role in the initial step of post-mitotic NPC reassembly. ELYS binds to the chromosome surface via its AT-hook domain (aa. 1971-1983) (18), which recruits early assembling Nups such as the Nup107-160 complex and Nups98 and 153. The AT-hook domain is located within the ELYS-C1 fragment examined in this study. Therefore, it is plausible that the C-terminal domain of ELYS functions as a platform for the successive assembly of FG-Nups via hydrophobic interactions among IDRs. In contrast, the structured N-terminal domain interacts directly with a helical domain of Nup120 (human Nup160) in the Y-complex (26). Therefore, it seems likely that the structured N-terminal domain of ELYS acts as a stereospecific binding site for the scaffold Y-complex, whereas the non-structured C-terminal region binds to the chromosome surface and initiates the ordered assembly of FG-Nups (Figure 4I).

Considering the facilitator function of ELYS and Nup35 in conjunction with the self-assembling characteristics of FG-Nups, it seems likely that reconstruction of the functional NPC during mitosis is driven by both promiscuous interactions among IDRs and stereospecific interactions among scaffold Nups, which are linked by unique Nup IDRs such as ELYS and Nup35N (Figure 4I). ELYS initiates the reassembly process by binding to the chromosome surface via its AT-hook domain. ELYS then recruits both scaffold Nups (Nups in the Y-complex), primarily via stereospecific interactions involving the N-terminus, and IDRs of early assembling FG-Nups (Nups153 and 98), primarily via promiscuous interactions involving the C-terminal IDR. The middle stage–assembling Nups58 and 62 cannot enter this IDR-rich phase at this stage. Nup35 eventually assembles into the IDR-rich phase formed at the ELYS C-terminus and then recruits the middle-to late-stage assembling Nups58 and 62. Simultaneously, the N-terminus of Nup35 appears to interact with the scaffold via the Nup93 complex (4) to link the FG-rich IDR phase to the entire scaffold. In the later stages, cytoplasmic Nups (Nups214 and 358) in addition to other Nups (Tpr, etc.) associate, potentially inducing further rearrangements of the IDR phase to establish the functional selective barrier within the NPC. It could be intriguing to investigate how phosphorylation/dephosphorylation of Nups (such as the Nup107-160 complex (27), Nup35 (28), and Nup98 (29) is involved in the regulation of Nup reassembly.

## Materials and Methods

### Amino acid sequence analyses

The disordered propensity was analyzed by DisProt server (http://disorder.compbio.iupui.edu/metapredictor.php) with a PONDR-fit algorithm. The net charge and the charge distribution were analyzed by a protein calculator (INNOVAGEN, https://pepcalc.com/protein-calculator.php), and EMBOSS explorer (http://www.bioinformatics.nl/cgi-bin/emboss/charge?_pref_hide_optional=1), respectively. The hydrophobicity was analyzed by ProtScale server in ExPASy (https://web.expasy.org/protscale/) by using Kyte & Doolittle method. COILS server in ExPASy (https://embnet.vital-it.ch/software/COILS_form.html) was used for coiled-coil analysis.

### DNA construction and protein purification

cDNA fragments encoding human Nup50FG (aa. 1–145), rat Nup62FG (aa. 1–268), human Nup98FG (aa. 214–480), human Nup153FG (aa. 1190–1464), human Nup214FG (aa. 1786–2090), human Nup214-C (aa. 1251–1550), human Nup35N (aa. 1–160), human Tpr-IDR1 (aa. 1451–1689), and human Tpr-IDR2 (aa. 1690–1937) were amplified by PCR from the full-length cDNA and sub-cloned into pET28-a (+) or pET28-b (+) (Novagen). Human Nup58FG-N (aa. 1–250), -C (aa. 438–599), human ELYS-IDR-M (aa. 1310–1559), human ELYS-IDR-C1 (aa. 1851–2033), and human ELYS-IDR-C2 (aa. 2034–2275) were artificially synthesized with the codon usage optimized for bacterial expression (Thermo Fisher Scientific), and sub-cloned into pET28-a (+). For IDRs lacking endogenous cysteine residue, a single cysteine was added to the carboxyl-terminal end of the fragment either by PCR or during artificial synthesis.

The expression of His_6_-tagged IDR-fragment in *Escherichia coli* cells (BL21-CodonPlus(DE3)-RIL, Agilent Technologies, Wilmington, DE) was induced by 0.5 mM IPTG at 37°C for 3 hours. The cells were harvested by centrifugation (5,000x g, 4°C, 15 min), and dissolved in Denaturing Buffer (8 M urea, 100 mM NaPO_4_, 10 mM Tris-HCl, 5 mM 2-mercaptoethanol (2-Me) (Nacalai Tesque, Kyoto, Japan), 10 mM imidazole, pH 8.0) at 4°C for 16 hours with a gentle agitation. An insoluble fraction was separated by centrifugation (10,000x g, 4°C, 30 min), and the supernatant was then incubated with Ni-NTA agarose beads (QIAGEN) at 4°C for 2 hours or at room temperature for 30 min. The beads were washed for several times with Denaturing Buffer containing 20 or 25 mM imidazole. His-tagged protein was eluted by 200 or 300 mM imidazole in Denaturing Buffer.

Purified protein was dialyzed against 0.1% (v/v) trifluoroacetic acid (TFA) (Nacalai Tesque) with 2 mM 2-Me at 4°C for 3 hours, and subsequently against 0.1% (v/v) TFA with 2 mM 2-Me at 4°C for ~20 hours. It was finally dialyzed against 0.05% (v/v) TFA, without 2-Me at 4°C for 2-3 hours, lyophilized (FDU-2200, EYELA), and stored at 4°C until use. Purified protein for CD measurement (Nup58CC (aa. 256-381) and Nup58FG-N) was first dialyzed against a buffer containing 100 mM L-arginine (50 mM KPO_4_, 100 mM L-arginine, 150 mM NaCl, 2 mM 2-Me), then against 50 mM L-arginine, and finally against 50 mM HEPES (pH 7.4) with 100 mM NaCl and 2 mM 2-Me at 4°C for 1 day, and stored −80°C by flash freezing with liquid nitrogen. The final protein concentration was determined by using Coomassie brilliant blue (CBB) staining, with bovine serum albumin (BSA) as a standard.

### Protein labeling

Purified IDR-Nups’ fragments were labeled by utilizing maleimide reaction with artificially added cysteine in the carboxyl terminal of each IDR-fragment or endogenous cysteine for some Nups (Nups214 and Nup58FG-C). Using ATTO dyes (ATTO390, 488, and 610, ATTO-TEC) dissolved in anhydrous DMSO (Nacalai Tesque), the appropriate volume of the ATTO solution was added in the protein stock buffer (final, 10 mM) before dissolving the lyophilized proteins. To stop the maleimide reaction, final 5 mM of dithiothreitol (DTT, Nacalai Tesque) was added into the denaturing protein stock solution and incubated at 4°C for more than 12 hours.

### In vitro self-assemble particle assay

Lyophilized proteins were dissolved in a small volume (~50 μL) of protein stock buffer (2 M Guanidine hydrochloride, 100 mM Tris-HCl, 10 mM HEPES, pH 8.0) and stored at 4°C for 1 day. This stock solution (≈1200 μM) was quickly diluted (>100 times) into the assay buffer (50 mM HEPES, 100 mM NaCl, pH 7.4) containing varying concentrations of crowding agents, salt, or 1,6-hexanediol (Nacalai Tesque), after removing aggregates by centrifugation (5 min at 10,000 x g). The final protein concentration was ≈ 2 μM for the client proteins and ≈3 μM for the host protein.

### Microscopic observation, image processing and analysis

Protein solution containing self-assembled particles was applied to a 96-well glass bottom plate (IWAKI, Shizuoka, Japan) or a custom-made flow chamber. In the mixing assay described in Figure 3, images were taken 5-10 min after the dilution of mixing host and client proteins in the chamber. In this time range, no significant change in the particle formation was observed. In the sequential addition of two different clients, 1st client protein was first added to the host particle (ELYS-C1). After 300 sec, fluorescence signal was captured by microscope, and 2nd client protein was added. After another 300 sec, the fluorescence signal of the 2nd client was observed under the microscope and subjected to the quantification. Microscopic observation was performed using FV3000 confocal laser-scanning microscope (OLYMPUS, Tokyo, Japan) using 60 × NA1.42 objective lens (PLAP0ON60XO) with oil immersion. For the total particle mass analysis, 10 images were randomly taken in 2x zoom field of 60 × objective lens. The excitation wavelength against ATTO390, 488, and 610 was 405 nm, 488 nm, and 564 nm, respectively. Sampling speed was 8.0 μs per pixel.

For *in vivo* live cell imaging, EGFP-fused Nups62 or 153 was co-expressed with mPlum (Clontech)-fused histone H3 in HeLa cells cultured in DMEM (SIGMA) supplemented with 10% FBS (Hyclone). Before the microscopic observations, the culture medium was replaced with phenol red-free medium. The stage-top chamber (Tokai-hit, Shizuoka, Japan) was filled with moisture using distilled water and 5 % CO_2_, and was maintained at 37°C. Images were captured every 1 min between metaphase and early G1 phase. Signal intensity at chromosome rim was measured as described below.

Microscopic images were exported as raw tiff 8-bit images (512×512 pixels) and analyzed with particle analyzer plugin in FIJI macro (https://fiji.sc/). When analyzing self-assembled particles, the sum of the RawIntDen (Raw Integrated Density) of each particle per image (total particle mass) was measured. For obtaining the partition coefficient (PC) of a client protein, a threshold for the particle size (>1 μm in Feret’s diameter) was applied to eliminate non-self-assembled aggregations. The ratio of the mean intensity of the client protein in the particle ([I]_particle_) over that of the bulk solution ([I]bulk) was calculated after the background subtraction and shown as common logarithm values. For the sequential partitioning assay, the threshold for the particles’ size was set between 5 and 15 μm in Feret’s diameter to avoid both small and large particles.

For analyzing EGFP signal at the chromosome surface in mitotic HeLa cells, the chromosome rim was detected in mPlum-H3 image and defined as a region of interest (ROI), which is then transferred to the image of EGFP channel. EGFP signal intensity in the ROI was quantified, subtracted by that of extrachromosomal space (background), and normalized by the value in interphase. Obtained data were arranged in Excel (Microsoft) and R version 3.4.1.

### FRAP analysis

The self-assembled particle was formed by ATTO488-labeled IDR-Nups as described above. A half or a center of the particle was bleached by a 488 nm laser at the maximum output for 4 sec. After bleaching, the time lapse observation was continued every 10 sec. Signal intensity of the bleached region was quantified manually using the FIJI and the obtained values were processed as mentioned above, and shown as a relative intensity to the signal intensity before bleaching.

### CD spectra measurement

The CD spectra of purified IDR-Nups and a coiled-coil protein (Nup58CC) were measured using a CD spectrophotometer (J-805, JASCO, Tokyo, Japan) with a 0.1 cm quartz cuvette. Data were acquired every 0.1 nm between 200 and 260 nm. Samples were prepared in assay buffer (50 mM HEPES pH 7.4, 100 mM NaCl, 2 mM 2-Me). The final protein concentration of Nup58FG-N, Nup58CC, Tpr-IDR1, and Tpr-IDR2 were 0.2, 0.2, 0.3, and 0.3 mg/mL, respectively.

### Nup35 knockdown

Small interference RNAs for Nup35 and control (luciferase) were purchased from Thermo (Silencer Select siRNA #29400 for Nup35 and Stealth™ RNAi 12935-146 for Luciferase). It was introduced into HeLa cells by Lipofectamine RNAi MAX (Invitrogen) by following the manufacturer’s protocol.

### Statistical analysis

To analyze the statistics for all of obtained data in this study, Welch two sample *t* test was used for normally distributed data, and Wilcoxon Rank-Sum test was used for non-normally distributed data. A Shapiro-Wilk normality test was used to determine whether data were distributed normally. All of statistical tests were done by using R version 3.4.1.

## Author Contributions

H.A.K. planned the project, designed the experiments, performed all experiments, and wrote the paper. S.H.Y. planned the project, designed the experiments, performed live cell imaging, and wrote the paper.

## Acknowledgements

We thank Prof. T. Haraguchi (human Nup98), Prof. K. Ullman (rat Nup62 and human Nup153), Prof. J. Ellenberg (human Nup214) for providing cDNAs encoding nucleoporins. We also thank S. Dodo for technical support, especially recombinant protein preparation and members of Yoshimura lab for discussions. This study was supported financially by the Funding Program for Next Generation World-leading Researchers (S.H.Y., No. LS076), and a Grant-in-Aid for Scientific Research (B) (S.H.Y., No. 21370054) from the Japan Society for the Promotion of Science (JSPS).

---

Table S1 Amino acid compositions and characteristics of IDR-Nups used in this study

Table S2 Partition coefficient of client protein against host protein in the assay shown in Figure 3. Data are summarized and represented as mean ± SD from more than two independent experiments. n indicates the number of analyzed particles. NA: not assigned.

Figure S1 Self-assembly of IDR-Nups

(A) Disorder probability of Nups used in this study. The results of PONDR-fit algorithm prediction are shown against amino acid position.

(B-E and J-M) DIC and fluorescence images of self-assembled particles formed by IDRs from FG-Nups (B-E) and non-FG-Nups (J-M) in different concentrations of PEG3350 (left). The total particle mass (see Experimental Procedures) was measured and summarized in the right panel. Data are presented as mean ± SEM, from three independent experiments. (*: p<0.05, **: p<0.005, ***: p<0.0005). P-values were obtained by appropriate statistical tests (see Experimental Procedures). (F-I, N and O) DIC and fluorescence images obtained from non-self-assembling Nups in the presence of 15% PEG3350. Scale bar: 10 μm.

Figure S2 Characterization of the self-assembling IDR-Nups

(A) Disorder probability of Tpr-IDRs. The result of PONDR-fit algorism prediction is shown.

(B) Co-line plot of the coiled-coil propensity (red) and the helix propensity (blue) of Tpr-IDRs.

(C) CD spectra of Tpr-IDR1 and Tpr-IDR2, Nup58FG-N and Nup58 (aa. 25-381). It was previously reported that Nup58FG-N is disordered and Nup58CC forms a stable coiled-coil (30).

(D-I) FRAP analysis of self-assembled particles (Nup98FG (D), Nup153FG (E), Nup214FG (F), Tpr-IDR1 (G), Nup35N (H), ELYS-C1 (I)). Fluorescence images before and after (0, 60, and 300 sec) bleaching (left panels), and signal intensity plot over time after the bleaching (right panel) are shown. Two different bleaching patterns (half; blue, centre; red) were tested. Data are represented as mean ± SD from three independent experiments.

Figure S3 Investigation of the driving force of self-assembly

(A and B) Fluorescence images of self-assembled particles in different concentrations of 1,6-hexanediol (A) and NaCl (B).

(C) Co-line plot of charge (red) and hydrophobicity (black) of ELYS-C1 against the amino acid position.

(D) Amino acid sequence of Nup35N fragment. Proline and glycine residue are labeled in green and red, respectively.

(E) Fluorescence images of non-colocalized particles shown as an asterisk (*) in Figure 3D. Scale bar: 2 μm.

Figure S4 Sequential assembly of IDR-Nups into a self-assembled particle of ELYS-C1

(A) DIC and fluorescence images of the host (ELYS-C1, blue) and the client proteins (green). Images were taken 3 min after the addition of the client. Scale bar: 20 μm.

(B) DIC and fluorescence images of the host (ELYS-C1, blue) and the 1st client (Nup35N or Nup98FG, red). The incorporation of the 1st client into the host particle was confirmed before the addition of the 2nd client protein. Scale bar: 20 μm.

(C) RNAi of Nup35. Small interference RNA for human Nup35 or luciferase was introduced into HeLa cells. The cell lysate was subjected to immunoblot analysis using anti-Nup35 and anti-β-actin antibodies. The position of Nup35 (~35 kDa) is indicated by an arrow.

## References

1. Maximiliano A. D’Angelo and Martin W. Hetzer. (2008) Structure, dynamics and function of nuclear pore complexes. Trends Cell Biol. 18, 456–466

2. Elad, N., Maimon, T., Frenkiel-Krispin, D., Lim, R. Y., and Medalia, O. (2009) Structural analysis of the nuclear pore complex by integrated approaches. Curr. Opin. Struct. Biol. 19, 226–232

3. von Appen, A., Kosinski, J., Sparks, L., Ori, A., DiGuilio, A. L., Vollmer, B., Mackmull, M.-T., Banterle, N., Parca, L., Kastritis, P., Buczak, K., Mosalaganti, S., Hagen, W., Andres-Pons, A., Lemke, E. a., Bork, P., Antonin, W., Glavy, J. S., Bui, K. H., and Beck, M. (2015) In situ structural analysis of the human nuclear pore complex. Nature 526, 140–143

4. Fischer, J., Teimer, R., Amlacher, S., Kunze, R., and Hurt, E. (2015) Linker Nups connect the nuclear pore complex inner ring with the outer ring and transport channel. Nat. Struct. Mol. Biol. 22, 774–781

5. Ullman, K. S. and Powers, M. A. (2015) Locking down the core of the pore. Science (80-.). 350, 33–34

6. Konishi, H. A., Asai, S., Watanabe, T. M., and Yoshimura, S. H. (2017) In vivo analysis of protein crowding within the nuclear pore complex in interphase and mitosis. Sci. Rep. 7, 5709

7. Rabut, G., Doye, V., and Ellenberg, J. (2004) Mapping the dynamic organization of the nuclear pore complex inside single living cells. Nat. Cell Biol. 6, 1114–1121

8. Denning, D. P., Patel, S. S., Uversky, V., Fink, A. L., and Rexach, M. (2003) Disorder in the nuclear pore complex: the FG repeat regions of nucleoporins are natively unfolded. Proc. Natl. Acad. Sci. U. S. A. 100, 2450–2455

9. Mohr, D., Frey, S., Fischer, T., Güttler, T., and Görlich, D. (2009) Characterisation of the passive permeability barrier of nuclear pore complexes. EMBO J. 28, 2541–2553

10. Frey, S. and Görlich, D. (2007) A saturated FG-repeat hydrogel can reproduce the permeability properties of nuclear pore complexes. Cell 130, 512–523

11. Patel, S. S., Belmont, B. J., Sante, J. M., and Rexach, M. F. (2007) Natively unfolded nucleoporins gate protein diffusion across the nuclear pore complex. Cell 129, 83–96

12. Frey, S., Richter, R. P., and Görlich, D. (2006) FG-rich repeats of nuclear pore proteins form a three-dimensional meshwork with hydrogel-like properties. Science 314, 815–817

13. Schmidt, H. B. and Görlich, D. (2015) Nup98 FG domains from diverse species spontaneously phase-separate into particles with nuclear pore-like permselectivity. Elife 4, 1–30

14. Dultz, E., Zanin, E., Wurzenberger, C., Braun, M., Rabut, G., Sironi, L., and Ellenberg, J. (2008) Systematic kinetic analysis of mitotic dis- and reassembly of the nuclear pore in living cells. J. Cell Biol. 180, 857–865

15. Otsuka, S., Szymborska, A., and Ellenberg, J. (2014) Imaging the assembly, structure, and function of the nuclear pore inside cells. Methods Cell Biol. 122, 219–238

16. Woodruff, J. B., Wueseke, O., Viscardi, V., Mahamid, J., Ochoa, S. D., Bunkenborg, J., Widlund, P. O., Pozniakovsky, A., Zanin, E., Bahmanyar, S., Zinke, A., Hong, S. H., Decker, M., Baumeister, W., Andersen, J. S., Oegema, K., and Hyman, A. a. (2015) Centrosomes. Regulated assembly of a supramolecular centrosome scaffold in vitro. Science 348, 808–812

17. Iwashita, K., Handa, A., and Shiraki, K. (2018) Coacervates and coaggregates: Liquid–liquid and liquid–solid phase transitions by native and unfolded protein complexes. Int. J. Biol. Macromol. 120, 10–18

18. Gillespie, P. J., Khoudoli, G. a, Stewart, G., Swedlow, J. R., and Blow, J. J. (2007) ELYS/MEL-28 chromatin association coordinates nuclear pore complex assembly and replication licensing. Curr. Biol. 17, 1657–1662

19. Otsuka, S. and Ellenberg, J. (2017) Mechanisms of nuclear pore complex assembly - two different ways of building one molecular machine. FEBS Lett. 592, 475–488

20. Kim, S. J., Fernandez-Martinez, J., Nudelman, I., Shi, Y., Zhang, W., Raveh, B., Herricks, T., Slaughter, B. D., Hogan, J. A., Upla, P., Chemmama, I. E., Pellarin, R., Echeverria, I., Shivaraju, M., Chaudhury, A. S., Wang, J., Williams, R., Unruh, J. R., Greenberg, C. H., Jacobs, E. Y., Yu, Z., de la Cruz, M. J., Mironska, R., Stokes, D. L., Aitchison, J. D., Jarrold, M. F., Gerton, J. L., Ludtke, S. J., Akey, C. W., Chait, B. T., Sali, A., and Rout, M. P. (2018) Integrative structure and functional anatomy of a nuclear pore complex. Nature 555, 475–482

21. Chug, H., Trakhanov, S., Hülsmann, B. B., Pleiner, T., and Görlich, D. (2015) Crystal structure of the metazoan Nup62•Nup58•Nup54 nucleoporin complex. Science 350, 106–110

22. Li, P., Banjade, S., Cheng, H.-C., Kim, S., Chen, B., Guo, L., Llaguno, M., Hollingsworth, J. V, King, D. S., Banani, S. F., Russo, P. S., Jiang, Q.-X., Nixon, B. T., and Rosen, M. K. (2012) Phase transitions in the assembly of multivalent signalling proteins. Nature 483, 336–340

23. Piela, L., Némethy, G., and Scheraga, H. A. (1987) Proline-induced constraints in α-helices. Biopolymers 26, 1587–1600

24. O’Neil, K. T. and DeGrado, W. F. (1990) A thermodynamic scale for the helix-forming tendencies of the commonly occurring amino acids. Science 250, 646–651

25. Rauscher, S., Baud, S., Miao, M., Keeley, F. W., and Pomès, R. (2006) Proline and Glycine Control Protein Self-Organization into Elastomeric or Amyloid Fibrils. Structure 14, 1667–1676

26. Bilokapic, S. and Schwartz, T. U. (2012) Molecular basis for Nup37 and ELY5/ELYS recruitment to the nuclear pore complex. Proc. Natl. Acad. Sci. 109, 15241–15246

27. Glavy, J. S., Krutchinsky, A. N., Cristea, I. M., Berke, I. C., Boehmer, T., Blobel, G., and Chait, B. T. (2007) Cell-cycle-dependent phosphorylation of the nuclear pore Nup107-160 subcomplex. Proc. Natl. Acad. Sci. U. S. A. 104, 3811–3816

28. Linder, M. I., Köhler, M., Boersema, P., Weberruss, M., Wandke, C., Marino, J., Ashiono, C., Picotti, P., Antonin, W., and Kutay, U. (2017) Mitotic Disassembly of Nuclear Pore Complexes Involves CDK1- and PLK1-Mediated Phosphorylation of Key Interconnecting Nucleoporins. Dev. Cell 43, 141–156

29. Laurell, E., Beck, K., Krupina, K., Theerthagiri, G., Bodenmiller, B., Horvath, P., Aebersold, R., Antonin, W., and Kutay, U. (2011) Phosphorylation of Nup98 by multiple kinases is crucial for NPC disassembly during mitotic entry. Cell 144, 539–550

30. Solmaz, S., Chauhan, R., Blobel, G., and Melčák, I. (2011) Molecular architecture of the transport channel of the nuclear pore complex. Cell 147, 592–602

