## Supplemental Figures for "Interactions between non-structured domains of FG- and non-FG-nucleoporins coordinate the ordered assembly of the nuclear pore complex in mitosis"

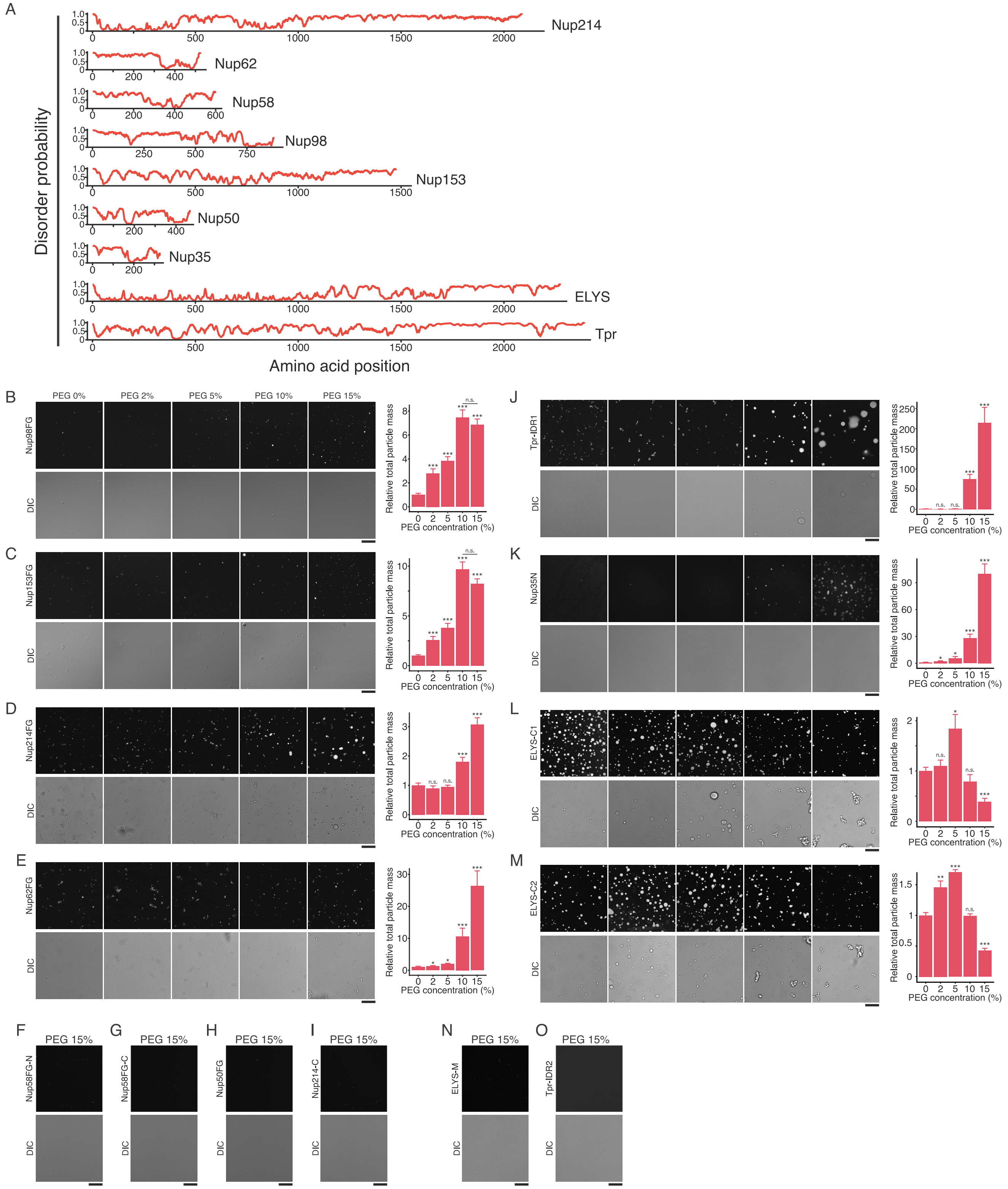


Konishi HA et al., Figure S1

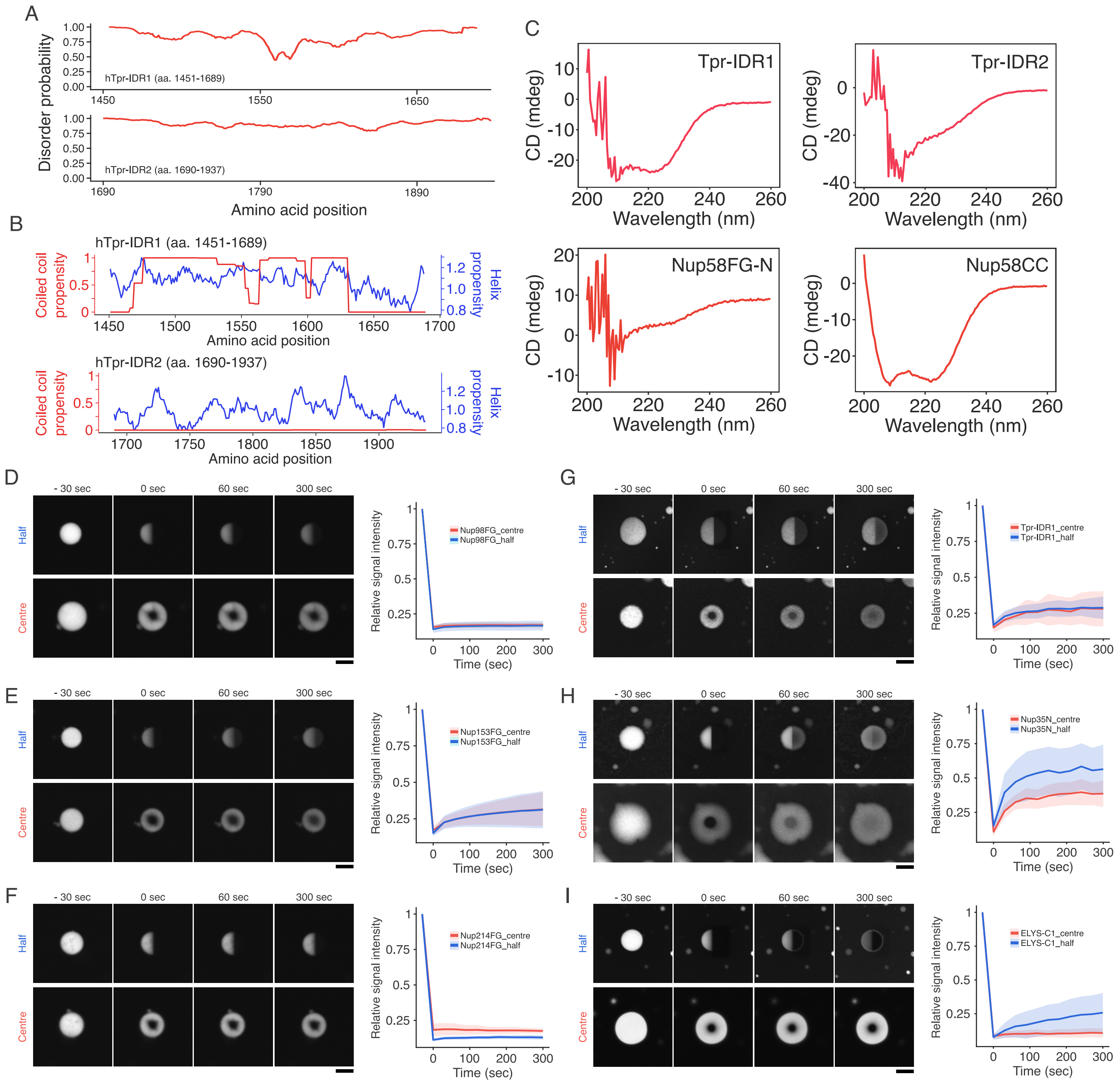


Konishi HA et al. Figure S2

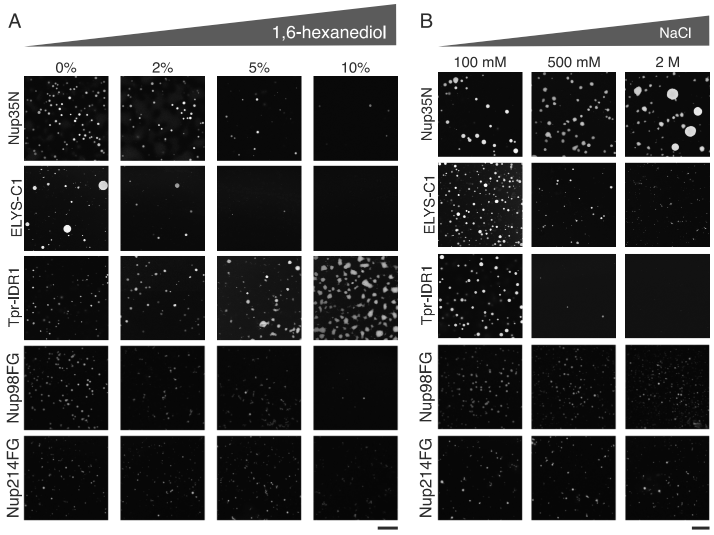


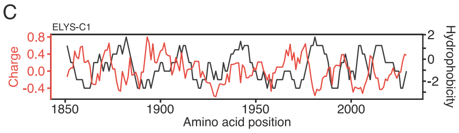


D

Nup35N (aa. 1-160)

1 MAAFAVEPQGPALGSEPMMLGSPTSPKPGVNAQFLPGFLM
41 GDLPAPVTPQPRSISGPSVGVMEMRSPLLAGGSPPQPVVP
81 AHKDKSGAPPVRSIYDDISSPGLGSTPLTSRRQPNISVMQ
121 SPLVGVTSTPGTGQSMFSPASIGQPRKTTLSPAQLDPFYT

28 prolines/160 aa (18%)

17 glycines/160 aa (11%)

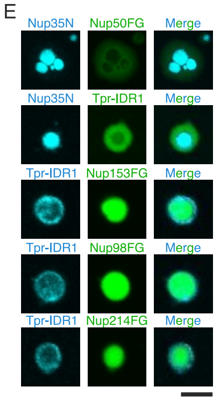


Konishi et al., Figure S3

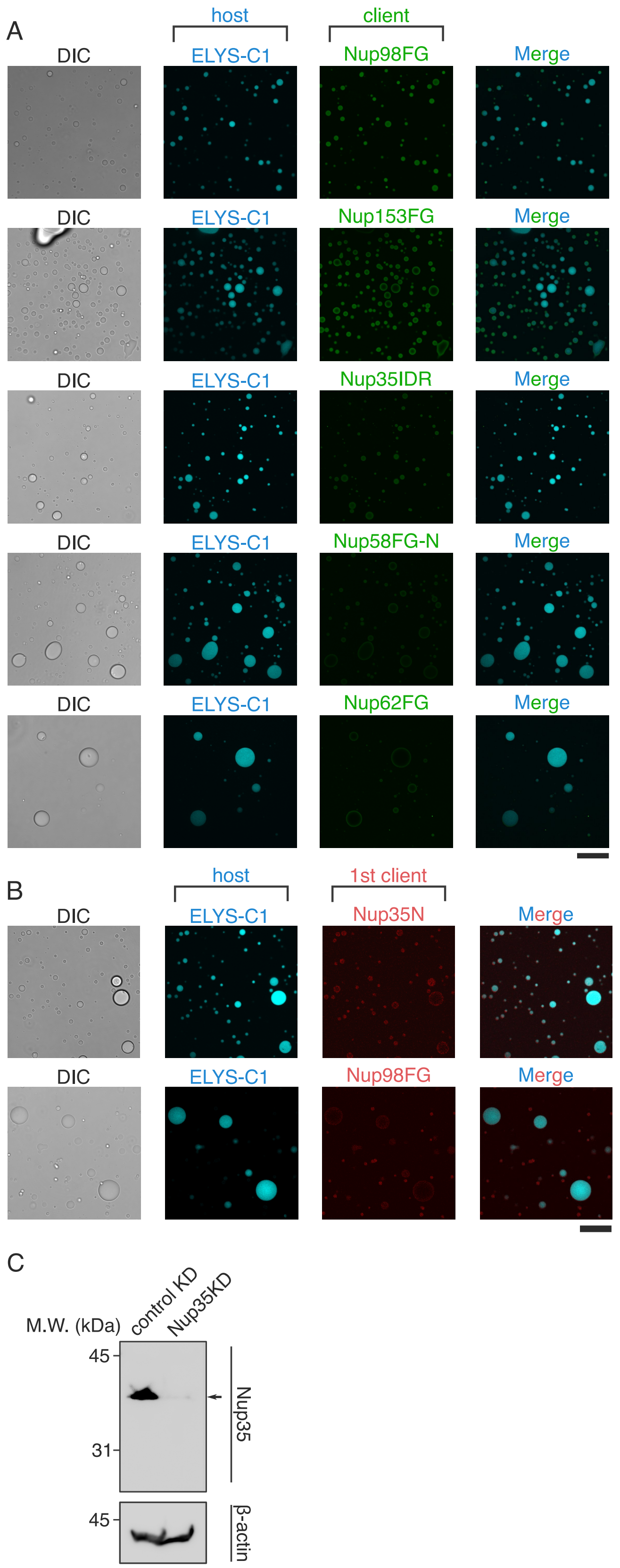


Konishi HA et al., Figure S4

| | | Name | Amino acid position | Molecular weight | Number of residues | Number of F, I, L, V, M, and P | Net charge | UniProt KB number |
|---|---|---|---|---|---|---|---|---|
| FG-Nups | Self-organized | hNup98FG | aa 214-480 | ≈ 26 kDa | 267 | 24 | 4 | P52948 |
| FG-Nups | Self-organized | hNup153FG | aa 1190-1464 | ≈ 28 kDa | 275 | 27 | 2 | P49790 |
| FG-Nups | Self-organized | hNup214FG | aa 1786-2090 | ≈ 30 kDa | 305 | 23 | 4.9 | P35658 |
| FG-Nups | Self-organized | rNup62FG | aa 1-268 | ≈ 28 kDa | 268 | 25 | 3 | P17955 |
| FG-Nups | Non-self-organized | hNup58FG-C | aa 438-599 | ≈ 17 kDa | 162 | 27 | 9.9 | Q9BVL2 |
| FG-Nups | Non-self-organized | hNup58FG-N | aa 1-250 | ≈ 25 kDa | 250 | 24 | 0 | Q9BVL2 |
| FG-Nups | Non-self-organized | hNup214-C | aa 1251-1550 | ≈ 31 kDa | 300 | 30 | -4.9 | P35658 |
| FG-Nups | Non-self-organized | hNup50FG | aa 1-145 | ≈ 16 kDa | 145 | 25 | 2 | Q9UKX7 |
| IDR-Nups (non-FG-Nups) | Self-organized | hNup35N | aa 1-160 | ≈ 17 kDa | 160 | 69 | 4.1 | Q8NFH5 |
| IDR-Nups (non-FG-Nups) | Self-organized | hELYS-C1 | aa 1851-2033 | ≈ 21 kDa | 183 | 47 | 18.1 | Q8WYP5 |
| IDR-Nups (non-FG-Nups) | Self-organized | hELYS-C2 | aa 2034-2275 | ≈ 27 kDa | 242 | 68 | 21.2 | Q8WYP5 |
| IDR-Nups (non-FG-Nups) | Self-organized | hTpr-IDR1 | aa 1451-1689 | ≈ 27 kDa | 239 | 53 | -5.5 | P12270 |
| IDR-Nups (non-FG-Nups) | Non-self-organized | hELYS-M | aa 1310-1559 | ≈ 27 kDa | 250 | 73 | -37.2 | Q8WYP5 |
| IDR-Nups (non-FG-Nups) | Non-self-organized | hTpr-IDR2 | aa 1690-1937 | ≈ 26 kDa | 248 | 73 | -25.8 | P12270 |

Konishi HA et al., Table S1

| host / client | BSA | Nup214FG -rich | Nup214FG -less | Nup62FG | Nup58FG-C | Nup58FG-N | Nup98FG | Nup153FG | Nup50FG |
|---|---|---|---|---|---|---|---|---|---|
| Nup214FG-rich | 0.51±0.12 (n=275) | 1.55±0.34 (n=1467) | 0.81±0.33 (n=409) | 1.71±0.25 (n=826) | 0.76±0.29 (n=1371) | 1.00±0.22 (n=641) | 1.48±0.26 (n=571) | 1.56±0.33 (n=778) | 0.33±0.10 (n=171) |
| Nup62FG | 0.67±0.17 (n=45) | 1.71±0.25 (n=826) | 1.30±0.17 (n=26) | 1.62±0.27 (n=480) | 1.05±0.25 (n=361) | 1.29±0.33 (n=274) | 1.58±0.32 (n=40) | 1.67±0.36 (n=158) | 0.60±0.13 (n=59) |
| Nup98FG | 0.19±0.09 (n=211) | 1.48±0.26 (n=571) | 1.24±0.27 (n=82) | 1.58±0.32 (n=40) | 0.73±0.26 (n=382) | 1.52±0.29 (n=205) | 1.41±0.26 (n=615) | 1.28±0.29 (n=340) | 0.82±0.20 (n=62) |
| Nup153FG | 0.46±0.11 (n=182) | 1.56±0.33 (n=778) | 0.94±0.24 (n=359) | 1.67±0.36 (n=158) | 1.28±0.33 (n=355) | 1.53±0.25 (n=237) | 1.28±0.29 (n=340) | 1.48±0.23 (n=505) | 0.58±0.18 (n=214) |

| host / client | BSA | ELYS-C1 | Nup153FG | Nup98FG | Nup35IDR | Nup58FG-N | Nup50FG | Nup62FG | Nup214FG -rich | Tpr-IDR1 |
|---|---|---|---|---|---|---|---|---|---|---|
| Tpr-IDR1 | 0.88±0.20 (n=59) | 1.02±0.14 (n=536) | NA | NA | NA | 0.99±0.21 (n=109) | 1.34±0.24 (n=205) | NA | NA | 1.17±0.20 (n=44) |
| Nup35N | 1.00±0.27 (n=66) | 1.28±0.30 (n=920) | 1.59±0.26 (n=417) | 1.53±0.29 (n=88) | 1.30±0.30 (n=538) | 1.28±0.24 (n=367) | NA | 1.62±0.24 (n=787) | 1.77±0.27 (n=1110) | NA |
| ELYS-C1 | 0.57±0.18 (n=215) | 1.57±0.29 (n=741) | 1.62±0.24 (n=704) | 1.23±0.27 (n=472) | 1.28±0.30 (n=920) | 1.13±0.22 (n=1951) | 0.57±0.15 (n=758) | 1.51±0.36 (n=1116) | 1.71±0.31 (n=1043) | 1.02±0.14 (n=536) |

Konishi HA et al., Table S2
